# Mechanism of assembly, activation and lysine selection by the SIN3B histone deacetylase complex

**DOI:** 10.1101/2023.01.26.525585

**Authors:** Mandy S.M. Wan, Reyhan Muhammad, Marios G. Koliopoulos, Theodoros I. Roumeliotis, Jyoti S. Choudhary, Claudio Alfieri

## Abstract

Histone deacetylase complexes remove histone lysine acetylation, a key post-translational modification that activates transcription at each gene. Although these complexes are drug targets and crucial regulators of organismal physiology, their structure and mechanisms of action are largely unclear. Here, we present the first structure of a complete human SIN3B histone deacetylase holo-complex with and without a substrate mimic. Remarkably, SIN3B encircles the deacetylase and contacts its allosteric basic patch thereby stimulating catalysis. A SIN3B loop inserts into the catalytic tunnel, rearranges to accommodate the acetyl-lysine moiety and stabilises the substrate for specific deacetylation, which is guided by a substrate receptor subunit. Our findings provide a model of specificity for a main transcriptional regulator conserved from yeast to human and a resource of protein-protein interactions for future drug designs.

## Introduction

Histone acetylation activates transcription by inhibiting nucleosome-DNA interactions thereby promoting chromatin decompaction, and by recruiting transcriptional co-activators that recognise this modification ^1–3^. This histone mark is “erased” by the activity of histone deacetylases (HDACs). Class I HDACs (HDAC1/2/3/8) represent the principle nuclear deacetylases that regulate transcription. HDAC1 and HDAC2 (HDAC1/2) are essential for cellular differentiation, development ^4–7^ and function as part of multi-subunit complexes ^8^. HDAC1/2 are Zn^2+^-dependent enzymes which catalyse the nucleophilic attack of a Zn^2+^-bound water molecule to a Zn^2+^-polarized N-∊-acetyl-lysine side-chain carbonyl. This generates a tetrahedral intermediate that collapses by protonation from a catalytic histidine residue, which generates acetate and lysine ^9^. Intriguingly, catalytic activity and substrate/gene specificity of HDAC1/2 in isolation is very poor or absent respectively ^8^. These physiological roles emerge when HDAC1/2 form specific complexes with co-repressor proteins such as the SIN3 proteins ^10–13^. How substrate specificity and catalytic activation is achieved in these holo-complexes is incompletely understood because we are lacking biochemical and structural information on the full-length version of these complexes, which are challenging to reconstitute and structurally analyse for their high intrinsic disorder.

In mammals, two highly similar proteins named SIN3A and SIN3B ^14^, take part in two distinct mammalian SIN3 complexes ^15,16^. These two complexes are homologous to the *Saccharomyces cerevisiae* Rpd3(L) and Rpd3(S) complexes respectively ^15,17–20^. Depletion of SIN3A causes loss of proliferative potential ^21–23^ and depletion of SIN3B inhibits the ability of proliferating cells to exit the cell cycle ^21,24–26^. Because of the ability of SIN3 complexes to regulate cell differentiation ^4–6^ and proliferation in cancer cells, they are drug targets ^27–30^. Consequently, several inhibitors have been developed and used in clinical trials ^31,32^, however success on this has been limited by the lack of specificity of these compounds towards the distinct HDAC1/2 complexes ^33^.

Understanding the assembly of different subunits, within the holo-SIN3 complexes is therefore critical to understand the mechanisms of substrate selection and activation in vivo and to design more specific and potent inhibitors for anti-cancer therapy. Here we report the first cryo-electron microscopy (cryo-EM) structure of the human SIN3B complex in apo form and bound to suberoanilide hydroxamic acid (SAHA) ^31,33^, which mimics an acetyl-lysine substrate. Our structural data combined with complementary cross-linking mass spectrometry (XL-MS) and biochemical assays show the molecular basis of complex assembly, complex-dependent stimulation of catalytic activity, and lysine selection in the context of an acetylated nucleosome substrate.

### Reconstitution of recombinant human SIN3B complex

It has been proposed that the human SIN3B complex consists of the SIN3 scaffold subunit, one catalytic core (i.e. HDAC1/2), and two chromatin targeting modules: the plant homeodomain (PHD) containing protein PHF12, and the MRG (MORF4-related gene) and chromo domain-containing protein named MORF4L1 (mortality factor 4-like 1) ^15,34^. However, clear evidence of direct protein:protein interaction among these subunits has been lacking. In order to address this, we reconstituted a recombinant version of the human SIN3B complex, composed of the subunits SIN3B isoform 2, HDAC2 (hereafter named HDAC for simplicity), PHF12 and MORF4L1 (Fig. 1a), expressed in the baculovirus/insect cell system. The resulting complex is stable in size-exclusion chromatography, stoichiometric, highly pure, and enzymatically active as it manifests HDAC activity (Supplementary Fig. 1a-b and d).

**Fig. 1.**
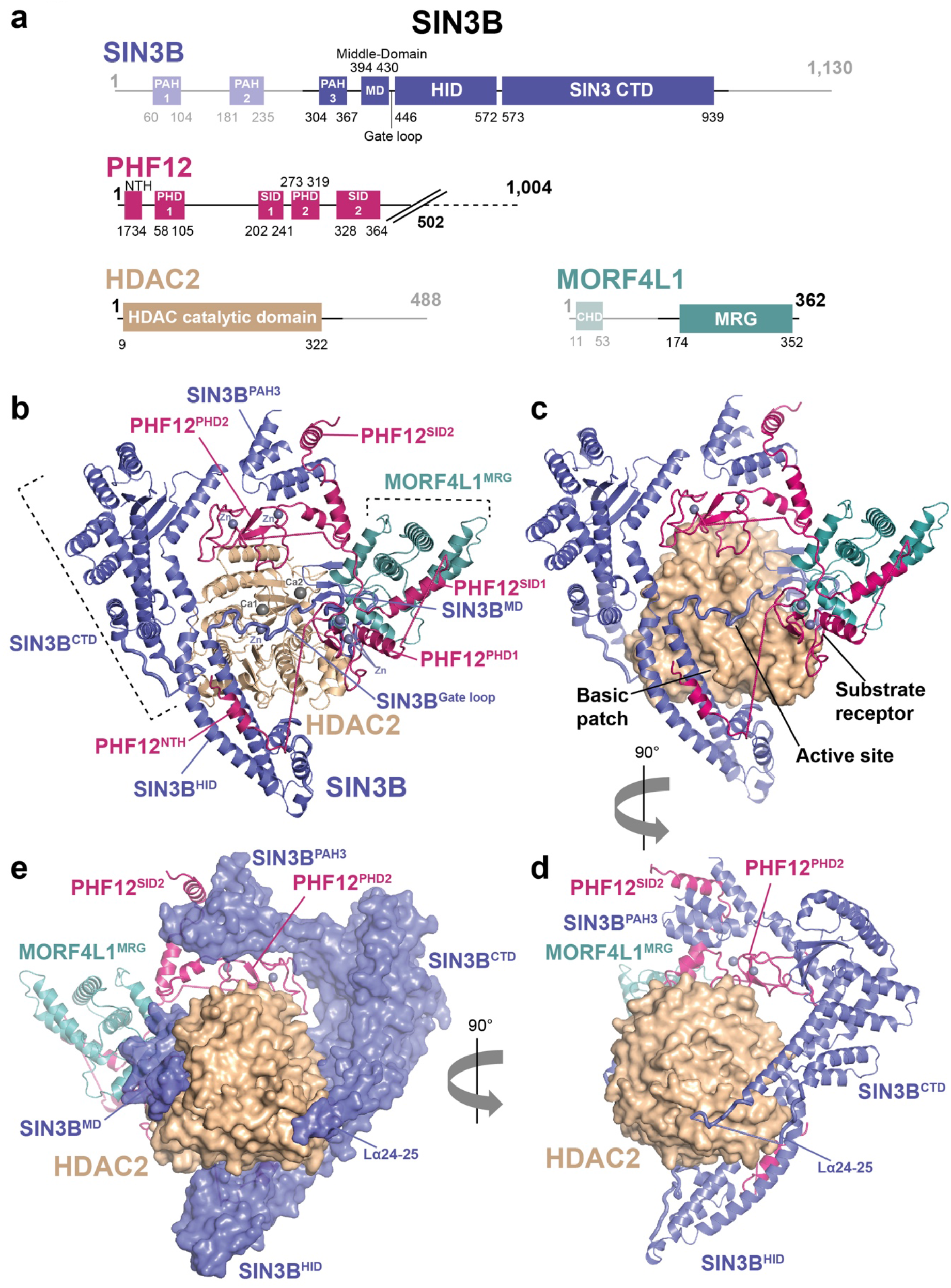
Overall architecture of the SIN3B apo complex. **a** Schematics showing domains and functional regions on the subunits of the SIN3B complex represented in linear form. Disordered parts of the complex are represented with fading colours. The PHF12^ΔC-terminus^ construct lacks the poorly conserved and unstructured C-terminus (as observed in our EM analysis) and it ends at residue 502. **b-d** Cartoon representations of three main views of the SIN3B complex showing the overall architecture of the complex. HDAC2 is represented as surface in **c,d** and **e**. SIN3B is represented as surface in **e**.

### Cryo-EM structure of the SIN3B complex

In order to characterise the stoichiometry and architecture of the SIN3B complex, as well as the molecular mechanism of HDAC activation by SIN3B, we pursued the structure determination of this complex by cryo-EM (see Methods and Supplementary Figs. 2, 3, 4 and Supplementary Table 1). 2D class-averages of this complex show V-shaped particles of ~150 Å size (Supplementary Figs. 2a and 3b). Ab-initio reconstruction of this complex followed by 3D-refinements and classifications allowed us to obtain a high-resolution map of the full-length SIN3B complex at 3.4 Å resolution and an additional map of the catalytic SIN3B core subcomplex (SIN3B^core^) at the resolution of 2.8 Å (Supplementary Figs. 2, 3, 4, 5 and Supplementary Movie 1). The SIN3B^core^ encompasses most of the SIN3B protein, including its third paired amphipathic α-helix (PAH3) domain, a previously unknown middle-domain (MD), the HDAC interacting domain (HID) domain, and a SIN3B C-terminal domain (CTD), which was previously defined ^10^ as PAH4 and highly conserved region (HCR) (Fig. 1a-b). Strikingly, SIN3B wraps around the catalytic HDAC module with its MD, HID, and one loop coming from the CTD (Fig.1a-e). Moreover, a loop (i.e. Gate loop) connecting MD and HID inserts into the catalytic site of the HDAC (Fig. 1a-c). SIN3B^core^ also includes PHF12, which mediates interactions with all the subunits of SIN3B. PHF12 includes one previously uncharacterised N-terminal helix (NTH), one PHD domain (PHD2), and one previously predicted SIN3 interacting domain ^35^ (SID2) (Fig. 1a-b). In the SIN3B complex, an additional histone recognition module sits on top of SIN3B^MD^ and HDAC, and includes the MRG domain of MORF4L1, which interacts with the PHF12^SID1^, and the first PHD domain of PHF12 (PHD1) (Fig. 1a-c, Supplementary Fig. 5a and Supplementary Movie 1).

**Fig. 2.**
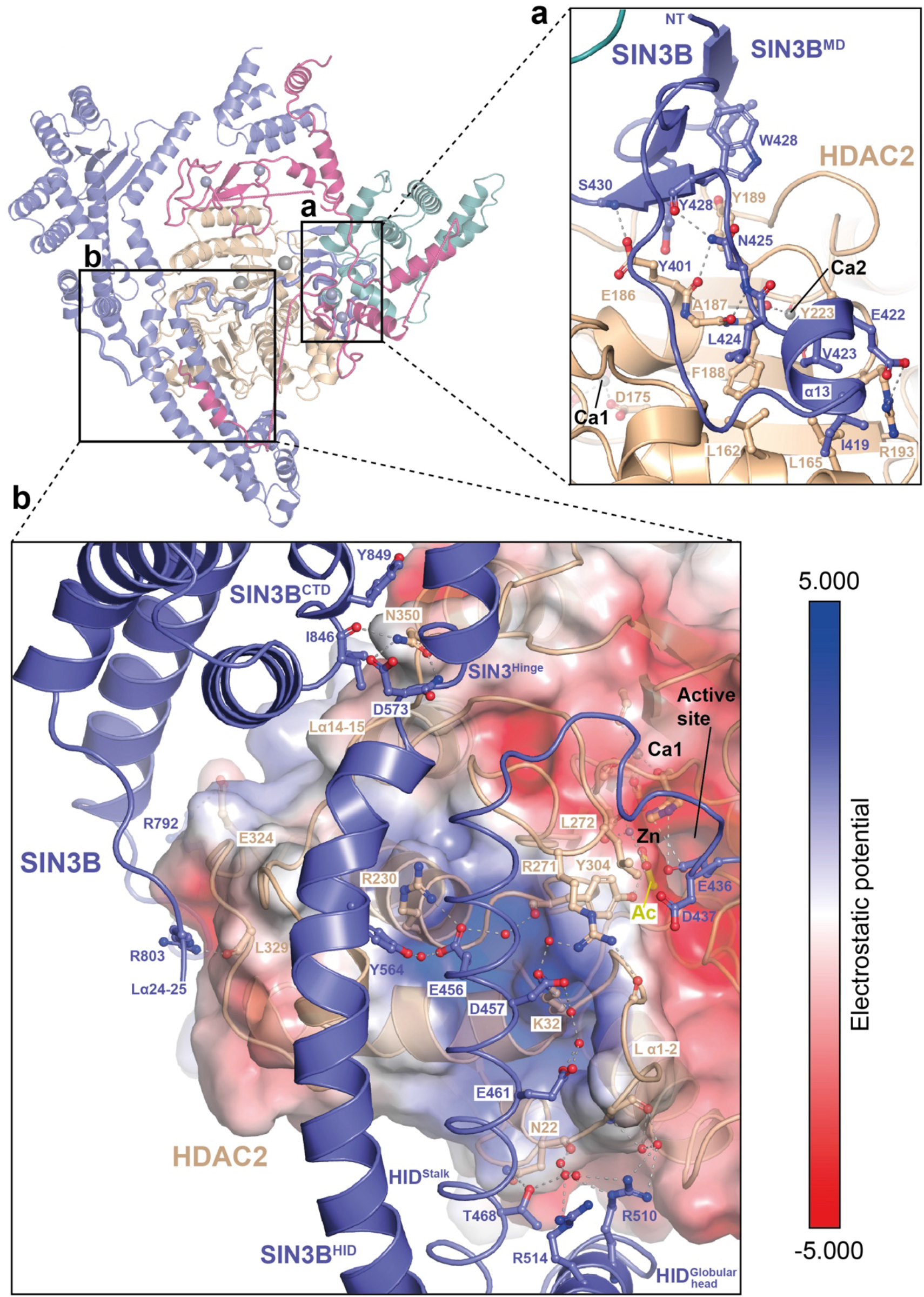
HDAC activation by SIN3B. **a** Close-up view on the interaction between the SIN3B middle domain (SIN3B^MD^) and the HDAC second calcium ion (Ca2) binding site. Residues involved in hydrophobic and backbone-mediated interactions are depicted. **b** Close-up view on the interaction between SIN3B HDAC interacting domain (HID) and the HDAC basic patch, which in other class I HDAC complex is involved in activation by Inositol phosphates (InsPs). The electrostatic surface potential (−/+5.000) is shown on HDAC. SIN3B^HID^ residues involved in interactions with HDAC2 are depicted. Most of these interactions are water mediated.

**Fig. 3.**
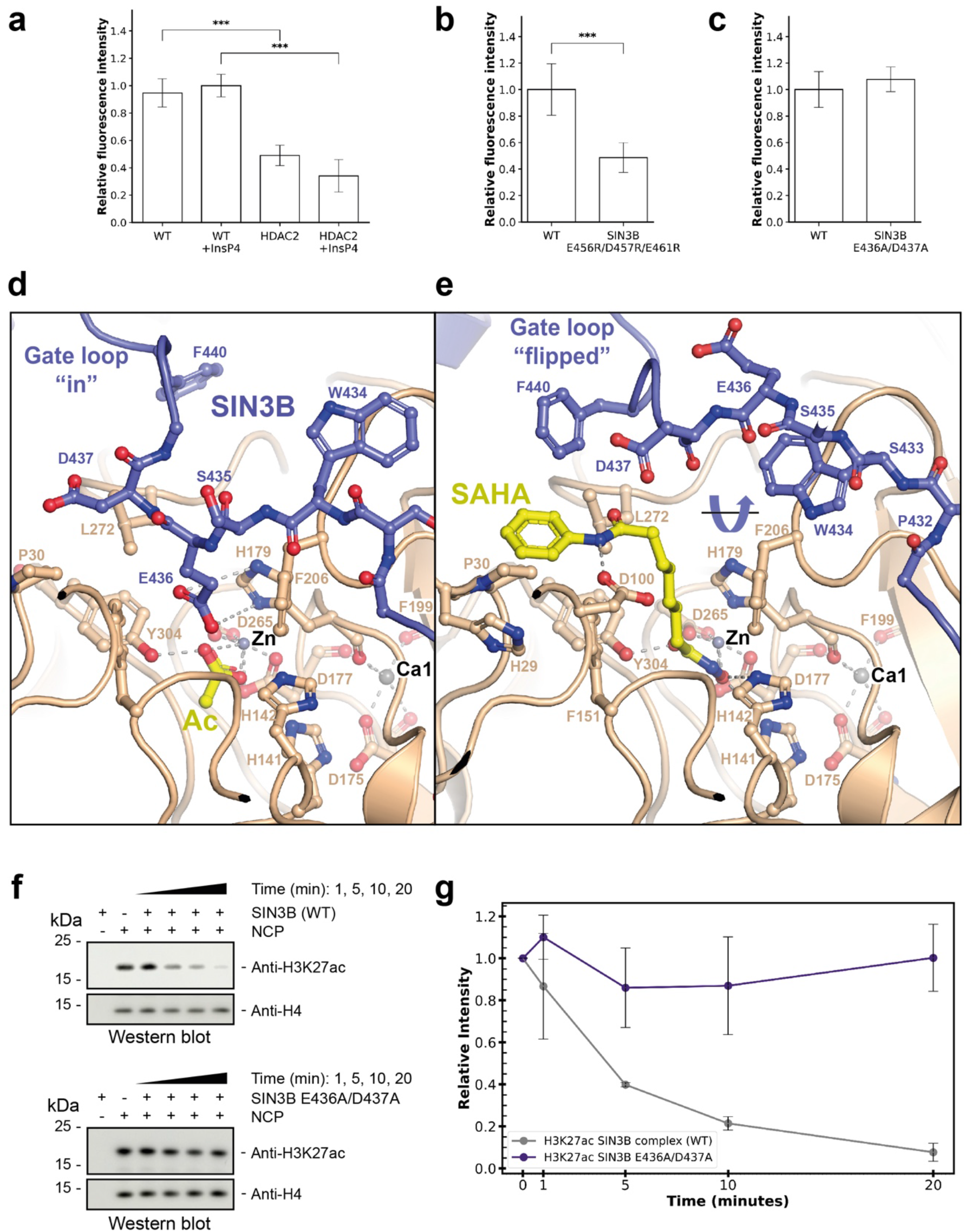
Structure of the SIN3B complex with the acetyl-lysine mimic SAHA. **a-c** HDAC activity assay performed with either HDAC alone, SIN3B complex and, SIN3B complex point mutants. Data are presented as mean values ± S.D. of independent experiments, n = 4 for (**a**), n = 6 (**b**) and n = 7 (**c**). ***p <0.0005 (one tailed unpaired t-test). **d-e** Close-up view on the catalytic core of either the SIN3B complex (**d**) or the SIN3B:SAHA complex cryo-EM structure determined here (**e**). The SIN3B^Gate loop^, which is displaced by SAHA binding is highlighted. **f-g** Deacetylation reactions by the SIN3B complex of nucleosomes with the acetyl moiety localised in lysine 27 (i.e. H3K27ac). The same experiment is also performed with a SIN3B Gate loop mutant (SIN3B^E436A/D437A^) complex. Quantifications from western blot data (one replicate is shown in **f**) is plotted in (**g**). Data are presented as mean values ± S.D. of independent experiments, n = 3.

**Fig. 4.**
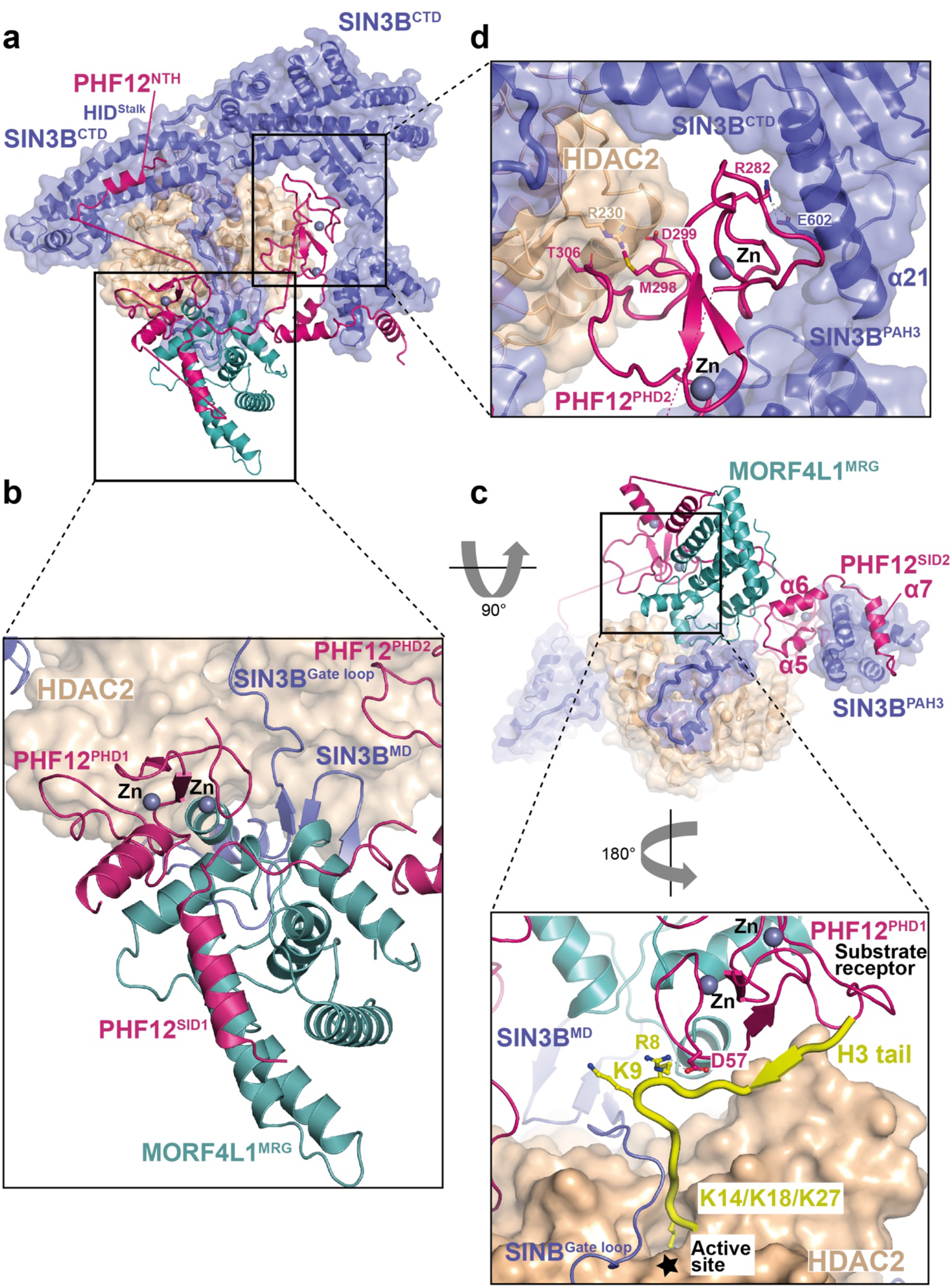
Substrate recognition by SIN3B. **a** Overview of the SIN3B complex structure represented as cartoon and transparent surface. PHF12 (magenta) is highlighted here and its PHD finger domains are shown in close up view in (**b**, PHD1) or (**d**, PHD2). **b** Close-up view on the PHF12 interactions with MORF4L1 MRG domain. PHF12^PHD1^ recognises a composite interface of MRG and the PHF12 SIN3B interaction domain 1 (SID1). **c** At the top, the structure in (**a**) is shown rotated 180 degrees to highlight the interactions of PHF12^SID2^ with SIN3B^PAH3^ and MORF4L1 MRG domain. At the bottom, it is shown the modelling of the H3 histone tail based on the structure of BHC80^PHD^:H3 superposed to our SIN3B structure (PHD1 as a reference) and based on our SIN3B:SAHA structure. Lysine residues on the H3 N-terminus are shown, K9 would be too far from the catalytic site, lysine residues more C-terminal to H3 residue 13 (i.e. K14, K18 and K27) would be able to enter the HDAC catalytic site. **d** Close-up view on the PHF12^PHD2^ interacting with SIN3B and HDAC2. Key residues involved in these interactions are depicted.

**Fig. 5.**
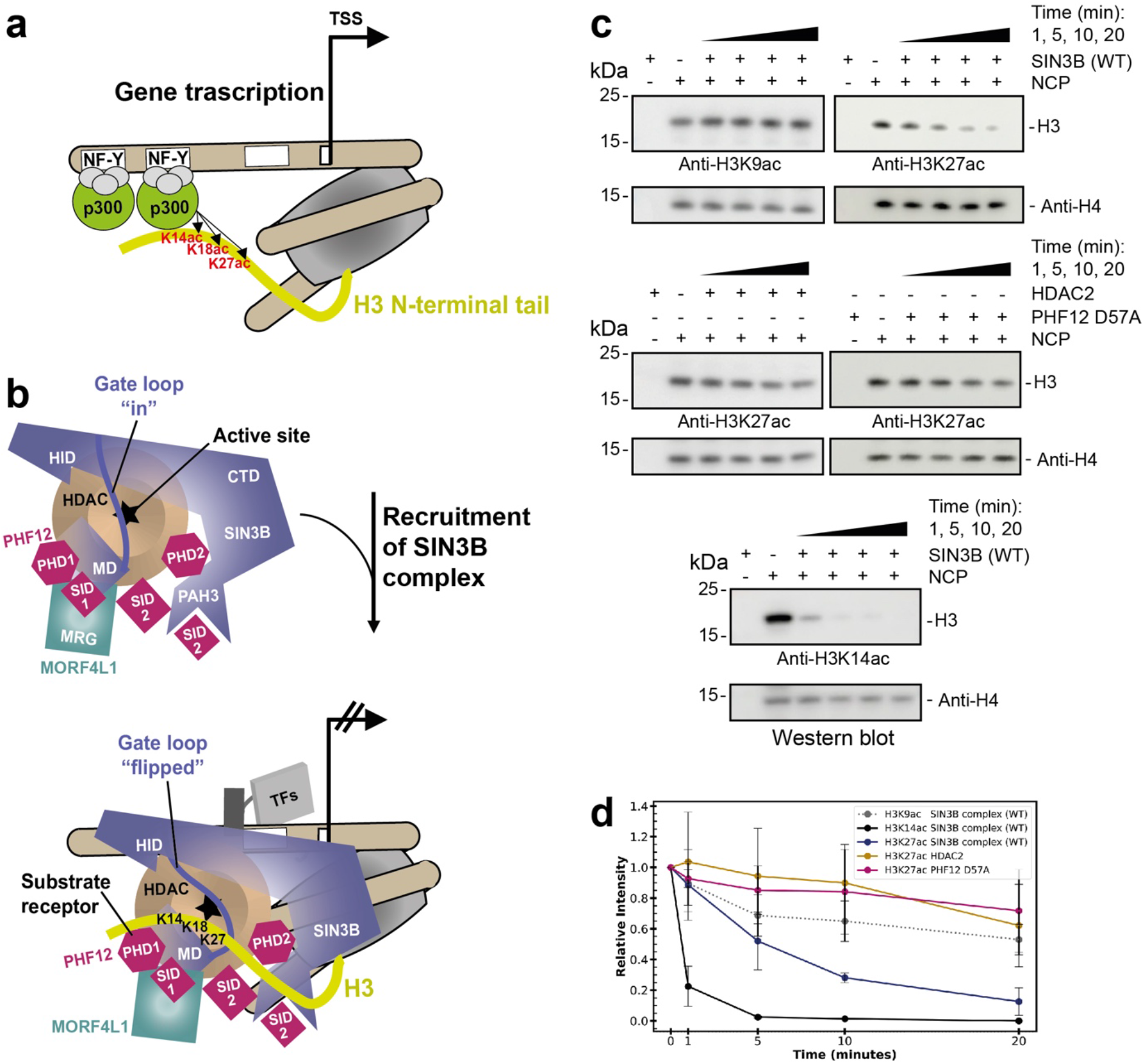
Model of SIN3B function at gene promoters. **a-b** Schematic cartoon illustrating a model for the function of SIN3B complex at target genes. SIN3B counteract p300-mediated acetylation. p300 binds the NF-Y sequence at target genes, whereas p300 can be recruited by various transcription factors (TFs) and their DNA-specific sequence (indicated with a white box). The HDAC active site is indicated with a star, the activating functions of the middle domain (MD) and HDAC interaction domain (HID) of SIN3B is indicated with a lightning symbol. The conformational change of the SIN3B^Gate loop^ upon histone H3 tail binding is indicated. H3K27/K18/K14 (and not H3K9) are main acetylation targets by p300 ^56,60^. **c-d** Deacetylation reactions by the SIN3B complex of nucleosomes with the acetyl moiety localised in different positions along the H3 histone tail (i.e. H3K9ac, H3K14ac and H3K27ac). The H3K27 deacetylation experiment is also performed with a SIN3B PHF12 PHD finger 1 mutant (SIN3B^PHF12D57A^) complex and with the HDAC2 apo enzyme. Quantifications from western blot data (performed in triplicate and one replicate is shown in **c**) is plotted in (**d**). Data are presented as mean values ± S.D. of independent experiments, n = 3.

In conclusion, SIN3B, HDAC, PHF12 and MORF4L1 interact with a 1:1:1:1 stoichiometry to form a V-shaped complex where the SIN3B^CTD^ and substrate recognition module form two arms at the top, the SIN3B^HID^ at the bottom, the HDAC is in the centre, and the SIN3B^PAH3^ connects the two arms in such a way that the HDAC is completely encircled by the SIN3B subunits (Fig. 1e and Supplementary Movie 1).

### SIN3B establishes multiple interactions with HDAC, which stimulate catalysis

The SIN3B interaction with HDAC is extensive and buries about one third of the HDAC surface area (i.e. 3566 Å^2^) (Supplementary Movie 1, Fig. 1b-d). The first remarkable interaction between SIN3B and HDAC involves the SIN3B^MD^, a short domain containing a beta-sheet and one short helix (α-13). The beta-sheet forms by the contribution of two beta-strands N-terminal to α-13 and one strand C-terminal of α-13 (Fig. 1a-c and e, Fig. 2a, and Supplementary Fig. 6). SIN3B^MD^ packs against the second calcium (Ca2) binding site of HDAC, which has been shown to be important for the activity of class I HDACs (Fig. 2a) ^9,36^. This interaction features mainly hydrophobic contacts involving SIN3B^MD^ beta sheet Tyr401 with HDAC Tyr189, and SIN3B α-13 Leu424 with HDAC Leu162 and Phe188. Moreover, SIN3B Asn425 and HDAC Glu186 are involved in side-chain-backbone interactions which stabilise the MD:HDAC interaction (Fig. 2a). Consistently, our XL-MS data on the SIN3B complex show several cross-links between the MD and residues near HDAC Tyr223, the backbone of which interacts with the Ca2 atom (Supplementary Fig. 7a, Supplementary Table 2 and Fig. 2a). In SIN3B isoform 1, Leu424 (Fig. 2a and Supplementary Fig. 6a) is replaced by one aspartate residue, which would interfere with the hydrophobic contacts between MD and HDAC. This provides a structural explanation to the results from previous HDAC activity assays showing that SIN3B isoform 2 is more active than the isoform 1 ^34^.

In conclusion, SIN3B interacts extensively with HDAC, among these interactions the SIN3B^MD^ contacts the Ca2 binding site of HDAC thereby stimulating catalytic activity.

### The SIN3B^HID^ directly contacts the HDAC basic patch for catalytic activation

It has been shown that in the context of NuRD ^37^, MiDAC ^12,33^, and SMRT ^38^ complexes a SANT domain of a co-repressor subunit stimulates the HDAC activity via binding of an inositol phosphate (InsP) which functions as a molecular “glue” between the SANT domain and the enzyme. Subsequent studies have shown that InsP contacts an allosteric basic patch formed by the HDAC α1 and α2, which influences residues in its catalytic site ^38,39^.

Strikingly, in our structure the SIN3B HID domain binds directly the allosteric basic patch of HDAC, preventing InsP binding (Fig. 2b). This interaction is dominated by a triad of SIN3B acidic residues coming from the stalk region of the HID i.e., Glu456, Asp457 and Glu461 (Fig. 2b and Supplementary Fig. 6a). Specifically, Glu456 and Asp457 interact with HDAC Lys32 and Arg271 via a network of hydrogen bonds involving water molecules. The latter residues are involved in stimulating HDAC activity in HDAC complexes regulated by InsP ^37,38,40^. Moreover, Glu456 interact with the backbone of the catalytic Tyr304, which as illustrated below is involved in substrate binding. These interactions are consistent with cross-links we found between SIN3B HID and the C-terminal portion of HDAC2 (Fig. 2b and Supplementary Fig. 7a). The SIN3B^HID^ acidic triad occupies a similar position as the InsP in the structure of SMRT:HDAC complex ^38^ (Supplementary Fig. 8a). Consequently, in the SIN3B complex, the acidic triad of SIN3B^HID^ would compete with InsP binding. Therefore, it would be expected that the InsP would not further stimulate the catalytic activity of the SIN3B complex. We tested this hypothesis experimentally and found that although the HDAC activity in the SIN3B complex is higher than the one of the HDAC apo enzyme, there is no further stimulation after addition of inositol-1,4,5,6-tetraphosphate (InsP4) (Fig. 3a). These results suggest that the SIN3B acidic triad functionally replaces the InsP cofactor in the SIN3B complex. In order to assess that this is the case, we generated a SIN3B complex where the acidic triad of SIN3B^HID^ is replaced by arginine residues. Indeed, we found that the HDAC activity of this charge-swap mutant is compromised (Fig. 3b).

The HID:HDAC interaction is also buttressed by the globular head of the HID that packs against the HDAC loop connecting α1 with α2 (L α1-2) (Fig. 2b). At the opposite side, the SIN3B hinge between HID and CTD form a pocket for HDAC loop α14-15 Asn350 (Fig. 2b). Further down, the loop α24-25 coming from the SIN3B^CTD^ binds HDAC in such a way that this region and the SIN3B^MD^ hug completely HDAC from opposite sides (Fig. 1d-e and 2b).

In conclusion, our structural data show that SIN3B contacts HDAC in several key surfaces thereby stimulating HDAC activity and that this stimulation is independent from the InsP4 co-factor but relies on direct protein-protein interactions established by the SIN3B co-repressor. This represent a general novel mechanism of Class I HDAC activation by co-repressor proteins.

### The SIN3B^Gate loop^ inserts into the HDAC catalytic tunnel

As detailed above, our cryo-EM structure of the SIN3B complex shows that a loop connecting MD and HID domains (Gate loop) inserts into the active site of the enzyme (Fig. 1a-c, Fig. 2b, Supplementary Fig. 5b and Supplementary Movie 1). SIN3B Glu436 enters the active site and its carboxyl group establishes hydrogen bonds with the catalytic HDAC His179, whereas the Cβ and Cγ of Glu436 makes non-polar contacts with HDAC Phe206 (Fig. 3d). The Cβ of the following SIN3B Asp437 makes non-polar contacts with the HDAC hydrophobic Leu272 and Pro30. Similarly to ref.^37^, we could also find an acetate molecule coordinating with the catalytic Zn cation and the catalytic Tyr304 (Fig. 3d). This suggests that our SIN3B apo structure represents a post-hydrolysis state of the deacetylation reaction where the protein substrate has been released and the gate loop has returned to a conformation in which it occludes the active site.

### Acetyl-lysine recognition by the SIN3B complex and regulation by the SIN3B^Gate loop^

In order to gain insights into the acetyl-lysine substrate recognition by the SIN3B complex and into the functional role of the SIN3B^Gate loop^, we determined the cryo-EM structure of the SIN3B complex with the inhibitor SAHA, which mimics a substrate acetyl-lysine and the peptide backbone of the substrate +1 residue ^31,33^ (Supplementary Fig. 4f-g, Supplementary Table 1, and Supplementary Movie 1, and Fig. 3e). Strikingly, the SIN3B^Gate loop^, which in SIN3B apo structure occludes the catalytic site, flips 180 degrees, thereby providing access for SAHA binding in the catalytic tunnel (Supplementary Fig. 5b and f, Fig. 3d and e). SAHA replaces SIN3B residues Glu436 and Asp437 and the acetate molecule, which were present in the catalytic site of the SIN3B-apo structure, and its binding mode is comparable with the previous crystal structure of the same drug with HDAC ^41^ where the hydroxamic acid moiety coordinates the HDAC zinc cation, establishes hydrogen bonds with His141-142 and the catalytic Tyr304, and the aliphatic chain participates in hydrophobic contacts with HDAC Phe206 and stacks with HDAC His179 (Fig. 3e). The amide NH group connecting the aliphatic chain and the phenyl ring performs a hydrogen bond with HDAC Asp100, thereby giving specificity to this interaction by mimicking the substrate backbone +1 residue (Fig. 3e).

Strikingly, in our structure, the phenyl ring of SAHA which mimics a substrate +1 residue, stacks between the SIN3B^Gate loop^ Phe440 and the HDAC His29 and Pro30, thereby suggesting that the SIN3B^Gate loop^ participates in stabilising the histone substrate in the active site. In order to experimentally assess the functional importance of the SIN3B^Gate loop^ in deacetylation of a genuine histone substrate we tested the ability of SIN3B to deacetylate a nucleosome carrying the histone H3 lysine 27 acetylation (H3K27ac) (Fig. 3f and g). Strikingly, mutating the conserved Glu436 and Asp437 to alanine, which is predicted to displace the SIN3B^Gate loop^ away from the HDAC active site, abolishes the ability of SIN3B to deacetylate H3K27 from a nucleosome (Supplementary Fig. 6a and Fig. 3f and g). Our results show that the SIN3B^Gate loop^ should be in proximity of the active site for establishing transient interactions with the histone tail substrate and ensure substrate-specific deacetylation. Importantly, the same Gate loop mutant has no effect on the SIN3B general HDAC activity over an acetyl-lysine mimic (Fig. 3c) showing that Glu436 and Asp437 play a role only in substrate-specific HDAC activity by the SIN3B complex.

In conclusion, we find that histone deacetylation by the SIN3B complex involves the joint contribution of the HDAC active site residues and the SIN3B^Gate loop^, which together position the histone substrate for deacetylation. This is the first reported example to our knowledge where a co-repressor protein within an HDAC complex regulates the specific HDAC activity by contacting directly the substrate. Conversely, it has been reported the presence of an auto-inhibitory loop in regulating the activity of the p300 acetyltransferase (HAT) ^42^. Intriguingly, the SIN3B^Gate loop^ is ubiquitylated ^43^, in the future it would be interesting to investigate if these post-translational modifications could influence the HDAC activity of SIN3B similarly as SUMOylation influences the activity of p300-related HATs ^44^.

### PHF12 stabilizes the SIN3B:HDAC complex and forms a histone recognition module

Our structure shows that PHF12 is key in defining the SIN3B architecture since it contacts each core subunit domain and it also forms a histone tail recognition module (Fig. 1 and 4). Firstly, PHF12^NTH^ binds the stalk region of the HID in a way which is reminiscent of the SIN3A^HID^:SDS3^SID^ interaction ^45^ (Fig. 4a and Supplementary Fig. 8b). PHF12^NTH^ and SDS3^SID^ bind SIN3^HID^ in a mutually exclusive manner as the N-terminal extended portion of the SDS3^SID^ would clash with the C-terminal portion of the PHF^NTH^, which is in an extended conformation (Fig. 1 and Supplementary Fig. 8b). The PHF12^NTH^:SIN3B^HID^ interaction is also supported by our XL-MS data (Supplementary Fig. 7a).

Strikingly, PHF12^PHD1^ recognises a composite interface formed by the MORF4L1 MRG domain interacting with the PHF12^SID1^ as also supported by our XL-MS data (Fig. 4b-c and Supplementary Fig. 7). Our structural data is agreeing with previous binding studies conducted by NMR showing that PHF12^PHD1^ binds to MORF4L1^MRG^ in a PHF12^SID1^-dependnent manner ^46^. Furthermore, our structural data explains why the yeast MORF4L1 homologue (i.e. Eaf3) dissociates from the SIN3B complex when the SID1 of Rco1 (i.e. yeast PHF12) is deleted ^47^. The MRG:SID1:PHD1 ternary assembly form the histone recognition module (or substrate receptor) of SIN3B. Within this structure the PHF12^PHD1^ is located on top of the active site of the HDAC, suggesting that this domain could recruit the histone tail substrate to the enzyme. In fact, the PHF12^PHD1^ has been shown to bind the unmodified end of histone H3 ^46^. Molecular modelling of an H3 histone tail substrate:SIN3B complex based on (i) the acetyl-lysine position shown by our SIN3B:SAHA structure, and (ii) the superposition of the crystal structure of the BHC80^PHD^:H3 ^48^ with the PHD1 in our structure, indicates that PHF12 is indeed able to present the histone H3 tail to the catalytic tunnel of the HDAC starting from Lys14 (Fig. 4c). Conversely, PHF12^PHD2^ is an atypical PHD domain at the sequence level and from previous biochemical data it is not able to bind histone tails on its own ^46^. Furthermore, in our structure this domain is wedged between the SIN3B^CTD^ and the HDAC thereby stabilising the SIN3B:HDAC interaction (Fig. 4d). This interaction is mainly specified by the Glu602 of the SIN3B α21 which protrudes out from the SIN3B^CTD^ and by the PHF12 Arg282, which interacts with the HDAC Arg230 and PHF12 Met298, Asp299 and Thr306 (Fig. 4d). Therefore, our structure demonstrates that PHF12^PHD2^ has a structural role rather than a substrate recruitment role. Consistently, modelling of a PHF12^PHD2^:H3 complex manifests clashes of H3 with SIN3B and with a loop preceding PHF^PHD2^ (PHF^pre-PHD2^) (Supplementary Figs. 6b and 8c).

In conclusion, our data establishes that PHD1 can deliver the histone H3 substrate to the HDAC active site where only lysine residues C-terminal to residue 13 can enter (i.e. H3K14, K18 and K27) (Fig. 4c-d and Supplementary Fig. 8c).

### A second SID in PHF12 binds the SIN3B PAH3 domain

Our structure shows that a second SID C-terminal to the PHF12^PHD2^ binds to the SIN3B^PAH3^ (Fig. 4c). Importantly, this is consistent with our XL-MS data (Supplementary Fig. 7a). The PHF12^SID2^ is reminiscent of the SIN3A^PAH3^:SAP30 interaction ^49^ since both these SIDs are tri-partite and form an extensive interaction with the PAH domain (Supplementary Fig. 9). In our structure, the C-terminal α7 from PHF12 inserts into the PAH cleft formed by four amphipathic helices by means of hydrophobic interactions by the PHF12 helical motif VDFLNRIH (Supplementary Figs. 6b and Fig. 4c). More N-terminally, PHF12 α6 is loosely attached to the SIN3B^PAH3^ contacting rather the MORF4L1^MRG^ (Fig. 4c). Conversely, PHF12 α5 together with the preceding PHF12^PHD2^ packs against the PAH3 domain at the opposite side of the PHF12 helix 7, thereby stabilising the interaction of PAH3 with both PHF12 and α21 from SIN3B^CTD^ (Fig. 4c and d). Therefore, our structural data indicates that the PHF12^SID2^ is another crucial component involved in SIN3B complex assembly and it is critical for firmly anchoring the histone recruiting protein PHF12 to SIN3B. Functional importance of the PHF^SID2^ in targeting SIN3B to chromatin has been demonstrated in several biochemical studies ^50,51^. Firstly, a C-terminal deletion including the SID2 region in the yeast PHF12 (i.e. Rco1) affects nucleosome binding by the SIN3B complex. Secondly, a construct including the SID2 region recruits a functional SIN3B complex in a luciferase reporter assay.

In conclusion, we find that PHF12 is a key scaffolding subunit of the SIN3B complex which contributes to complex assembly by contacting each core subunit domain, and it constitutes a substrate receptor by recruiting the H3 histone tail with its PHD1. The structural importance of this subunit that we observe is consistent with the strong phenotypes observed in the PHF12 knockout ^52^. We also find that PHF12 competes with the SIN3A-associated SDS3 and SAP30 therefore explaining why these proteins take part in the two distinct SIN3A and SIN3B complexes ^20^.

### Specific SIN3B deacetylation of p300 histone marks on the histone H3

SIN3B contributes to gene repression by counteracting the activity of the HAT p300 at target promoters ^53–60^ (Fig. 5a-b). Our structure is consistent with a specific role of the SIN3B complex in removing the p300 specific mark H3K27 instead of the more N-terminal acetylation H3K9 (Fig. 4c). In order to demonstrate this, we probed deacetylation of SIN3B on nucleosomes containing either H3K9ac or H3K27ac marks. As result, SIN3B can deacetylate nucleosomes containing either H3K27ac or H3K14ac, but is unable to deacetylate nucleosomes containing H3K9ac, which is not a p300-specific histone mark (Fig. 5c and d). Strikingly, deacetylation of H3K27 is compromised if using either the HDAC in apo form or a SIN3B complex carrying the SIN3B^PHF12D57A^ point mutant which is unable to bind the histone H3 tail (Fig. 4c and 5c). This demonstrates that the SIN3B complex prefers deacetylating H3K27 over H3K9 and that H3K27 deacetylation depends on the SIN3B HDAC holo-complex assembly and specifically on the PHD finger 1 of PHF12 within the substrate receptor of SIN3B as shown in our structure.

In summary, here we report the first high-resolution structure of a full-length human HDAC complex which includes catalytic site, co-repressor HDAC activating subunit, and histone recognition module. We also report the first structure of a SIN3-family HDAC complex. We find a novel mechanism of activation for the drug target HDAC1/2 by the corepressor subunit SIN3B, which contacts the HDAC allosteric basic patch directly and completely encircles all the HDAC cation binding sites. By providing the structure of SIN3B complex with a substrate mimic, we also show structural evidence that the co-repressor SIN3B establishes transient interactions with the histone substrate for ensuring specific deacetylation. Moreover, we show that the histone recognition module of SIN3B recruits the histone tail via a PHD finger at a specific distance from the active site. This dictates which lysine residues can be fed in the catalytic tunnel, showing a specificity for H3K27ac over H3K9ac. Furthermore, the interaction between SIN3B^PAH3^ and PHF12 that we describe is important because the PAH domains of SIN3 proteins are drug targets ^61,62^. These findings pave the way for a model of specificity of HDAC complexes, which is crucial in understanding their general role in gene expression and chromatin structure regulation. Finally, our structure provides a resource of protein-protein interactions of an HDAC complex, which rationalises an ensemble of previous cellular and biochemical data and will guide the design of more specific and effective anti-cancer drugs.

## Acknowledgments

We thank Chris Richardson for the IT support. We thank Fabienne Beuron for her support within the ICR EM facility. Stephen Hearnshaw for his help within the ICR biophysics facility. We thank Ruth Knight for helping with insect cell culture. We thank Alex Radzisheuskaya, Basil J. Greber and Vlad Pena for critically reading and commenting on this manuscript. We thank David Barford for discussions and for allowing CA to start this project in his laboratory with a Brenner postdoc price awarded to CA (Max Peruz Fund, Laboratory of Molecular Biology, Cambridge 2018). We thank Diamond for access and support of the cryo-EM facilities at the UK national electron Bio-Imaging Centre (eBIC), proposal BI21809-34, funded by the Wellcome Trust, MRC and BBSRC.

We thank Chris Richardson for the IT support. We thank Fabienne Beuron for her support within the ICR EM facility. Stephen Hearnshaw for his help within the ICR biophysics facility. We thank Ruth Knight for helping with insect cell culture. We thank Alex Radzisheuskaya, Basil J. Greber and Vlad Pena for critically reading and commenting on this manuscript. We thank David Barford for discussions and for allowing CA to start this project in his laboratory with his Brenner postdoc price (Max Peruz Fund, Laboratory of Molecular Biology, Cambridge 2018). We thank Diamond for access and support of the cryo-EM facilities at the UK national electron Bio-Imaging Centre (eBIC), proposal BI21809-34, funded by the Wellcome Trust, MRC and BBSRC. MGK and CA are supported by the Sir Henry Dale Fellowship 215458/Z/19/Z. MW and RM are supported by the Institute of Cancer Research (ICR), grant number allocated are GFR005X and GFR146X respectively. The work of TIR and JSC was funded by the CRUK Centre grant with reference number C309/A25144.

## Author contributions

MW cloned and reconstituted all the SIN3B constructs in this study. MW performed all the biochemical assays in this study. RM performed GraFIX, prepared cryo-EM grids and helped CA to screen them. CA coordinated the EM pipeline, screened the cryo-EM grids, collected cryo-EM data, and processed the cryo-EM data. CA performed structural model interpretation and building with inputs from MGK. MGK performed atomic coordinate model refinement. TIR and JSC performed and supervised the MS analysis of the SIN3B apo sample respectively. CA directed the project and designed experiments with MW and RM. CA prepared the manuscript and wrote the manuscript with the help of MW and RM.

## Data and materials availability

The structural coordinates have been deposited in the Protein Data Bank with the following accession numbers: PDB-ID 8C60, 8BPA, 8BPB and 8BPC; the EM data have been deposited in the Electron Microscopy Data Bank with the following accession numbers: EMD-16449, EMD-16147, EMD-16148 and EMD-16149.

The mass spectrometry proteomics data have been deposited to the ProteomeXchange Consortium via the PRIDE^63^ partner repository with the dataset identifier.

All the other data and materials used for the analysis are available from the corresponding author upon reasonable request. Source data are provided with this paper.

## Supplementary Information

Materials and Methods

Supplementary Fig.s 1 to 9

Supplementary Table 1

Supplementary References

Supplementary Movie 1

## Supplementary Information

### Materials and Methods

#### Protein expression and purification

Codon optimised cDNAs of the SIN3B complex (C-terminally StrepII-tagged SIN3B isoform 2 (full length, CTD encompassing residues 301-943, E436A/D437A and E456R/D457R/E461R mutants), N-terminally GST-tagged MORF4L1, PHF12 (full-length, residues 1-502 and D57A mutant), RBBP7, and HDAC2 were purchased from GeneArt (Thermo Fischer Scientific) or GenScript and cloned into MultiBac vectors (pFastBac1 or pACEBac1) for insect cell expression.

Baculoviruses of each subunit were produced in Sf9 cells then co-infected in High Five cells at a density of 1.5 x 106 cells ml-1 using Sf-900 III medium supplemented with penicillin/streptomycin. Cells were shaken at 27 °C for 48 hours then harvested and stored at −80°C until processing.

Cell pellets were resuspended in cold lysis buffer (50 mM HEPES pH8, 500 mM NaCl, 0.5 mM TCEP, 5% glycerol, 1 mM EDTA, 0.5 mM PMSF, 2 mM benzamidine, 5 U ml-1 benzonase (Sigma), cOmpleteTM mini EDTA-free protease inhibitors (Roche)), lysed by sonication, and centrifuged. The supernatant was filtered and loaded on a Strep-Tactin Superflow (30060, Qiagen) column then washed with 10 CV 500 mM NaCl wash buffer (containing 500 mM NaCl, 50 mM HEPES pH 8, 5% glycerol, 1 mM EDTA), followed by 4 CV 1 M NaCl wash buffer, followed by 10 CV 200 mM NaCl wash buffer before elution using 200 mM NaCl wash buffer supplemented with 2.5 mM d-Desthiobiotin. The eluted fractions were subsequently loaded onto a GSTrapTM HP column preequilibrated with 200 mM NaCl wash buffer, followed by elution with 10 mM reduced glutathione in 200 mM NaCl. Samples were concentrated and the StrepII and GST tags were removed by O/N incubation with 3C protease at 4°C. SEC was used for final polishing on a Superose 6 Increase 10/300 GL column (Cytiva) in buffer containing 20 mM HEPES pH 8, 150 mM NaCl, and 0.5 mM TCEP. Peak fractions were pooled, concentrated up to 2 μM, and used for further experiments or flash frozen in liquid nitrogen and stored at −80°C.

In order to express the four-subunits SIN3B complex also either RbBP4 or RbBP7 must have been included in the co-expression. Although this protein is not visualised in any of our cryo-EM reconstructions, presumably because of its flexible tethering to the SIN3B complex, it allowed solubility of SIN3B. We also did not find any difference in the cryo-EM reconstructions between the SIN3B^FL^ complex and the SIN3B complex where we deleted the C-terminus of PHF12 (i.e. PHF12^ΔC-terminus^) thereby supporting the idea that also this portion of PHF12 is disordered.

#### NCP167 assembly

The unmodified NCP167 and the NCP167 with the H3K9/K14/K27ac modification were purchased from ActiveMotif.

#### Histone deacetylation activity assay using the Boc-Lys(Ac)-AMC substrate

For HDAC assays using the fluorogenic substrate, Boc-Lys(Ac)-AMC^64^, 100 nM protein was incubated with 100 μM Boc-Lys(ac)-AMC substrate (Cayman Chemical) in the presence or absence of 100 μM InsP4 in pH 7.4 TBS buffer for 1 hour. The reaction was stopped using 200 μM SAHA inhibitor for 5 minutes and deacetylated substrates were digested with 50 mg/mL trypsin for 1 hour. Fluorescence was measured using a POLARstar Omega (BMG Labtech) plate reader at 355/460 nm. Incubation steps were carried out at 37°C with agitation. Unpaired t-tests were applied to detect differences between samples and p-value <0.05 was considered as statistically significant.

#### Histone deacetylation activity assay using acetylated nucleosomes

100 nM H3K9ac or H3K27ac nucleosomes (Active Motif) were incubated on ice with 100 nM protein in buffer containing 25 mM HEPES pH 7.5, 50 mM NaCl, and 0.1 mM TCEP. Reactions were stopped at various timepoints by quenching with gel loading buffer (NuPAGE LDS sample buffer, Life Technologies), loaded onto 4-12% SDS-PAGE gels and transferred onto PVDF membranes for western blotting using site-specific primary antibodies, H3K9ac, H3K27ac, H3K14ac and H4 (Cell Signalling #9649, #D5E4, Sigma-Aldrich #07-353, and Abcam #ab7311 respectively). Membranes were subsequently probed with goat-anti-rabbit-HRP conjugated secondary antibody (Abcam, #ab205718) then incubated with ECL for detection and quantified using ImageJ.

#### Gradient Fixation (GraFix)

A continuous gradient containing 20 mM HEPES pH 7.5, 150 mM NaCl, 0.5 mM TCEP, 10-30% sucrose, 0-0.1% glutaraldehyde was generated using Gradient Master (Biocomp). 100 μL of SIN3B complex at the concentration of 1.7 μM was applied on top of the gradient, and centrifuged at 165,100g for 16 hours at 4 °C using SW 60 Ti ultracentrifuge. Samples were manually fractionated by hand, and the crosslinking was quenched by adding Tris-HCl pH 7.5 to a final concentration of 35 mM. GraFix ^65^ fractions were analysed on SDS-PAGE, and fractions that contain protein band close to the size of Sin3B complex were pooled, and buffer exchanged to 20 mM Tris-HCl (pH 8.0), 50 mM NaCl, 0.5 mM TCEP. For Sin3B:SAHA complex, SIN3B was mixed with 50 μM SAHA, and incubated for 30 minutes at 4 °C prior to GraFix. GraFix solution and final buffer exchange solution were supplemented with 50 μM SAHA.

#### Cryo-EM grid preparation

Quantifoil R1.2/1.3 Cu 300 grids were previously glow discharged using Easiglow (Pelco) at 15 mA for 1 minute. 2 μl of 1 μM GraFix-treated SIN3B/SIN3B:SAHA was applied to grids, and blotted for 5 seconds before being plunged into liquid ethane. Grids were prepared at 4 °C and 100% humidity using a Vitrobot mark IV (Thermo Fisher).

#### Cryo-EM data collection and processing

##### SIN3B-apo complex

We collected 48,837 movie stacks in MRC format on a Titan KRIOS 300 kV cryo-TEM at eBIC (Diamond light source, BI21809-37 session), equipped with a K3 detector operated in super-resolution bin 2x mode, at a nominal magnification of 165 kx, which yielded a pixel size of 0.513 Å per pixel. These movies were live-processed during collection using the cryoSPARC live program^66^.

Movie stacks were frame aligned and binned 4 times giving a pixel size of 2.052 Å per pixel. 41,727 Images with resolution better than 6 Å and a total motion of 30 pixels (estimated during frame alignment) were selected for further processing. A blob picker was used for initial particle picking during the live processing, and for obtaining initial 2D class averages. Templates generated from the latter, were used for template picking and TOPAZ^67^ training and picking with cryoSPARC. Selected particles from 2D classifications performed with particles picked with the different methods explained above were pooled together and duplicated particles were removed by using the “remove duplicates” function implemented in cryoSPARC. This step removed doubly picked particles within 50 Å (shortest dimension of the SIN3B particles is ~50 Å). These particles were piped into the “ab initio model” function implemented in cryoSPARC. These particles were then imported in RELION-3.1.1^68^ by using pyem and the csparc2star.py script by Daniel Asarnow: Daniel Asarnow, Eugene Palovcak, & Yifan Cheng. (2019). asarnow/pyem: UCSF pyem v0.5 (v0.5). Zenodo. https://doi.org/10.5281/zenodo.3576630. Refinement, Bayesian polishing was performed as in^69^. Further map improvement was performed by performing 3D classification without alignment (T=30) with a soft mask on the SIN3B ^core^ subcomplex and performing Ctf refinement in RELION. This yielded a map at the resolution of 2.8 Å. The SIN3B complex map was obtained by an initial overall 3D classification with angular sampling of 7.5 degrees, followed by 3D classification focusing on the substrate recognition module including (SIN3B^PHD1^:MORF4L1^MRG^:PHF^SID1^ subcomplex).

##### SIN3B:SAHA complex

We collected 29,661 Movie stacks in MRC format on a Titan KRIOS 300 kV cryo-TEM at eBIC (Diamond light source, BI21809-41 session), equipped with a K3 detector operated in super-resolution bin 2x mode, at a nominal magnification of 165 kx, which yielded a pixel size of 0.508 Å per pixel. Processing of these data was performed similarly to the SIN3B^core^ complex except that two cycles of Bayesian polishing followed by 3D classification were performed. The first 3D classification was performed at T=20, the second at T=40.

#### Model building

The PDB 6XEC [https://www.wwpdb.org/pdb?id=pdb_00006xec] was used as initial template for building HDAC2 in the SIN3B-apo structure in Coot^70^. SIN3B was build de novo based on the excellent quality of our cryo-EM map, AlphaFold^71^ models for the individual SIN3B domains were used as initial templates. The PDB 2LKM [https://www.wwpdb.org/pdb?id=pdb_00002lkm] was used as initial template to model the MORF4L1^MRG^:PHF12^SID1^ subcomplex. PHF12NTH and PHD1 and PHD2 were build de novo based on the excellent quality of our cryo-EM map, AlphaFold models for the individual domains were used as initial templates. Structural model refinement was performed with PHENIX real-space refinement^72^ at the resolution of 2.8 Å for the SIN3B^core^ complexes and at 4.1 Å for the SIN3B complex. For building the SIN3B:SAHA structure, the structure of SIN3B was used as an initial template, and the SAHA ligand was fitted unambiguously.

#### Crosslinking mass spectrometry analysis

30 μL of freshly purified SIN3B complex at a concentration of 1.7 μM was incubated with 0.75 μL DSSO (disuccinimidyl sulfoxide) (A33545, Thermo Scientific) pre-dissolved in DMSO at a concentration of 100 mM. Sample was incubated on ice for 10 min followed by quenching with 2 μL 500 mM Tris pH8. The sample was analysed by mass spectrometry as follows. After the crosslinking reaction, triethylammonium bicarbonate buffer (TEAB) was added to the sample at a final concentration of 100 mM. Proteins were reduced and alkylated with 5 mM tris-2-carboxyethyl phosphine (TCEP) and 10 mM iodoacetamide (IAA) simultaneously for 60 min in dark and were digested overnight with trypsin at final concentration 50 ng/μL (Pierce). Sample was dried and peptides were fractionated with high-pH Reversed-Phase (RP) chromatography using the XBridge C18 column (2.1 x 150 mm, 3.5 μm, Waters) on a Dionex UltiMate 3000 HPLC system. Mobile phase A was 0.1% v/v ammonium hydroxide and mobile phase B was acetonitrile, 0.1% v/v ammonium hydroxide. The peptides were fractionated at 0.2 mL/min with the following gradient: 5 minutes at 5% B, up to 12% B in 3 min, for 32 min gradient to 35% B, gradient to 80% B in 5 min, isocratic for 5 minutes and re-equilibration to 5% B. Fractions were collected every 42 sec, SpeedVac dried and orthogonally pooled into 12 samples for MS analysis. LC-MS analysis was performed on the Dionex UltiMate 3000 UHPLC system coupled with the Orbitrap Lumos Mass Spectrometer (Thermo Scientific). Each peptide fraction was reconstituted in 30 μL 0.1% formic acid and 15 μL were loaded to the Acclaim PepMap 100, 100 μm × 2 cm C18, 5 μm trapping column at 10 μL/min flow rate of 0.1% formic acid loading buffer. Peptides were then subjected to a gradient elution on the Acclaim PepMap (75 μm × 50 cm, 2 μm, 100 Å) C18 capillary column connected to a stainless steel emitter with integrated liquid junction (cat# PSSELJ, MSWIL) on the EASY-Spray source at 45 °C. Mobile phase A was 0.1% formic acid and mobile phase B was 80% acetonitrile, 0.1% formic acid. The gradient separation method at flow rate 300 nL/min was as follows: for 95 min gradient from 5%-38% B, for 5 min up to 95% B, for 5 min isocratic at 95% B, re-equilibration to 5% B in 5 min, for 10 min isocratic at 5% B. Precursors between 375-1,600 m/z and charge equal or higher than +3 were selected at 120,000 resolution in the top speed mode in 3 sec and were isolated for stepped HCD fragmentation (collision energies % = 25, 30, 36) with quadrupole isolation width 1.6 Th, Orbitrap detection with 30,000 resolution and 54 ms Maximum Injection Time. Targeted MS precursors were dynamically excluded for further isolation and activation for 30 seconds with 10 ppm mass tolerance.

Identification of crosslinked peptides was performed in Proteome Discoverer 2.4 (Thermo) with the Xlinkx search engine in the MS2 mode for DSSO / +158.004 Da (K). Precursor and fragment mass tolerances were 10 ppm and 0.02 Da respectively with maximum 2 trypsin missed cleavages allowed. Carbamidomethyl at C was selected as static modification. Spectra were searched against a FASTA file containing Human reviewed UniProt entries including the O75182-2 isoform. Crosslinked peptides were filtered at FDR<0.01 using the Percolator node and target-decoy database search.

**Supplementary Fig. 1.**
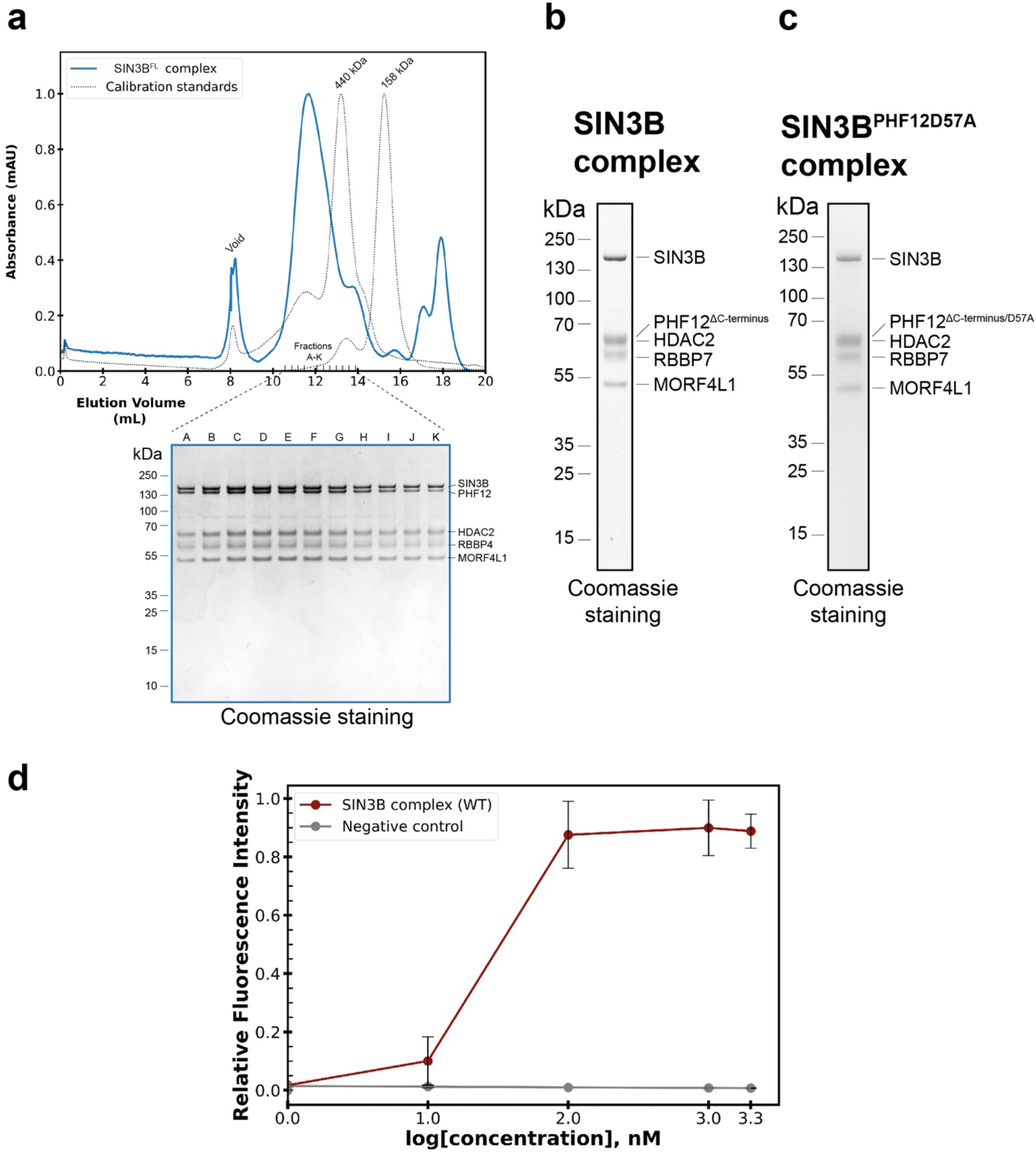
Biochemical reconstitutions of SIN3B complexes. **a** Size exclusion chromatography (SEC) chromatogram (Top) of the full length SIN3B complex (SIN3B^FL^). Calibration standards are illustrated for references of molecular weight. The SIN3B^FL^ complex elutes close to the 440 kDa molecular marker, the secondary peak eluting after the 158 kDa marker accounts for the 3C protease used to cleave the tags in SIN3B and MORF4L1 and the GST cleavage product. Coomassie stained gel of the eluted fractions (Bottom). **b-c** SDS PAGE of the protein complexes reconstituted in this study. **d** HDAC assay showing that the SIN3B complex is active. Data are presented as mean values ± S.D. of independent experiments, n = 3.

**Supplementary Fig. 2.**
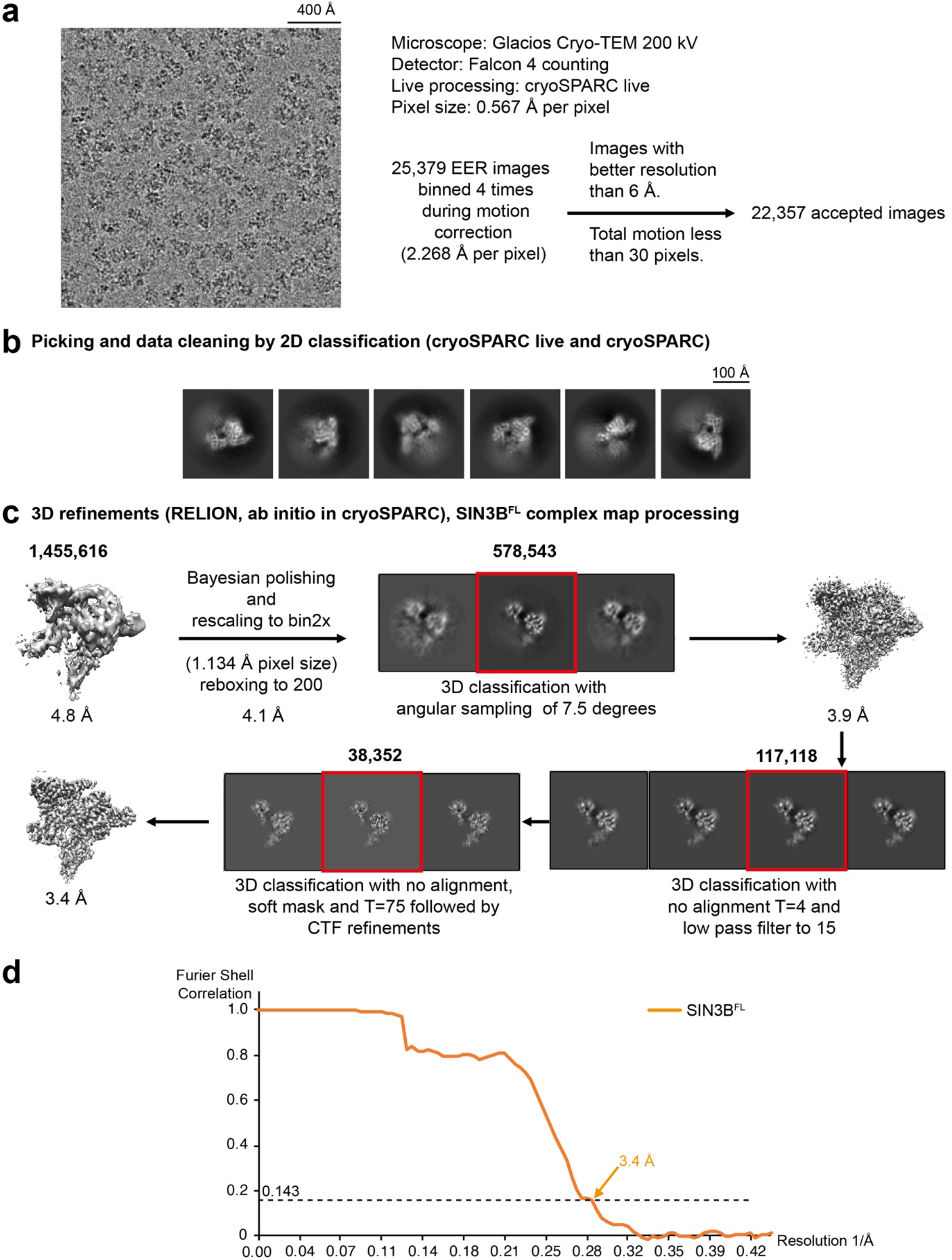
Cryo-EM analysis of the SIN3B full-length complex. Workflow showing a representative micrograph (low-pass filtered to 15 Å), the cryo-EM data collection parameters (**a**) and the single-particle analysis pipeline for the SIN3B full-length (SIN3B^FL^) complex (**b-c**). N. of particles at each classification step is indicated. More details are described in the main text and in the Methods section. **d** Fourier Shell Correlation (FSC) curve is shown.

**Supplementary Fig. 3.**
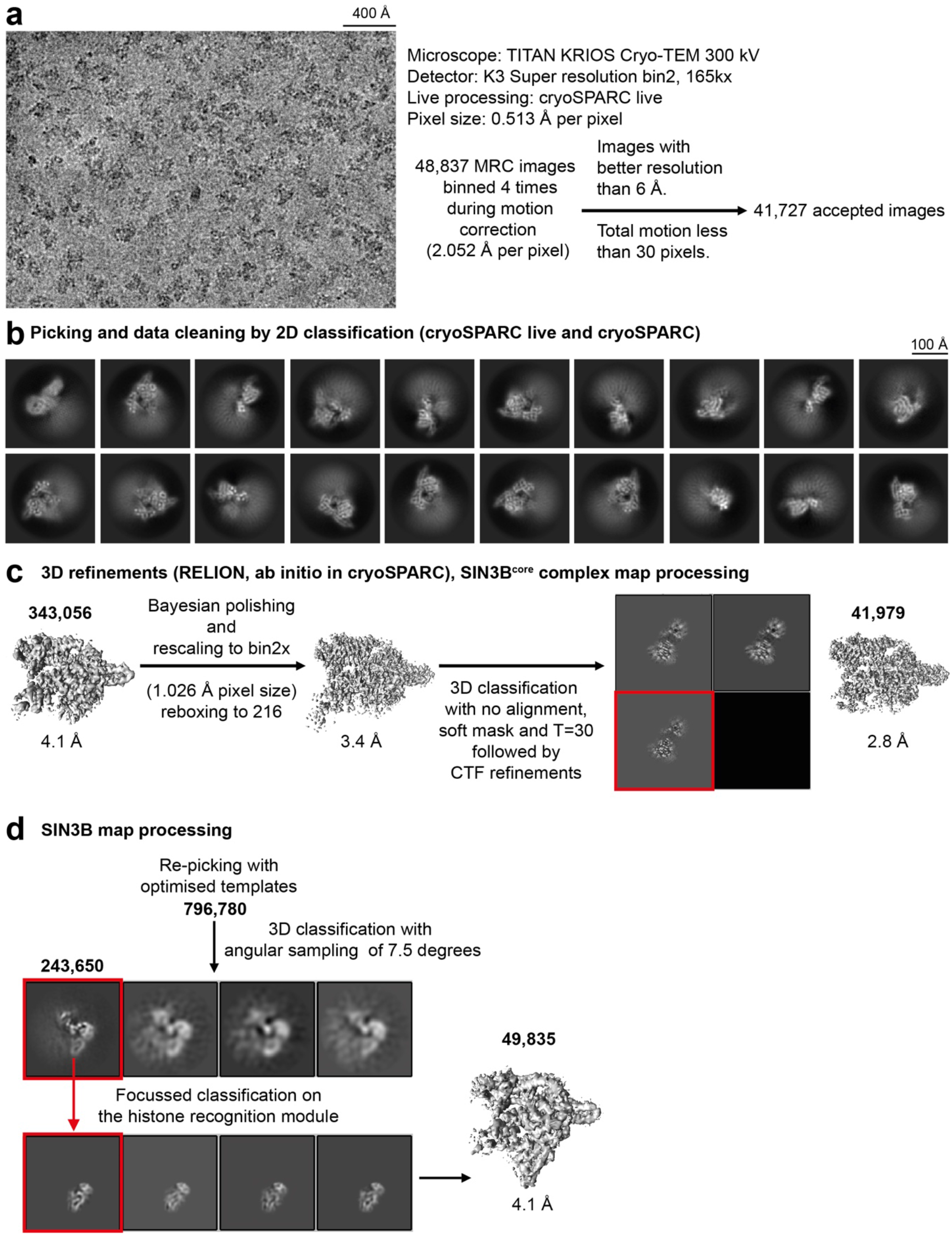
Cryo-EM analysis of the SIN3B complex. Workflow showing a representative micrograph (low-pass filtered to 15 Å), the cryo-EM data collection parameters (**a**) and the single-particle analysis pipeline (**b-d**). N. of particles at each classification step is indicated. More details are described in the Methods section.

**Supplementary Fig. 4.**
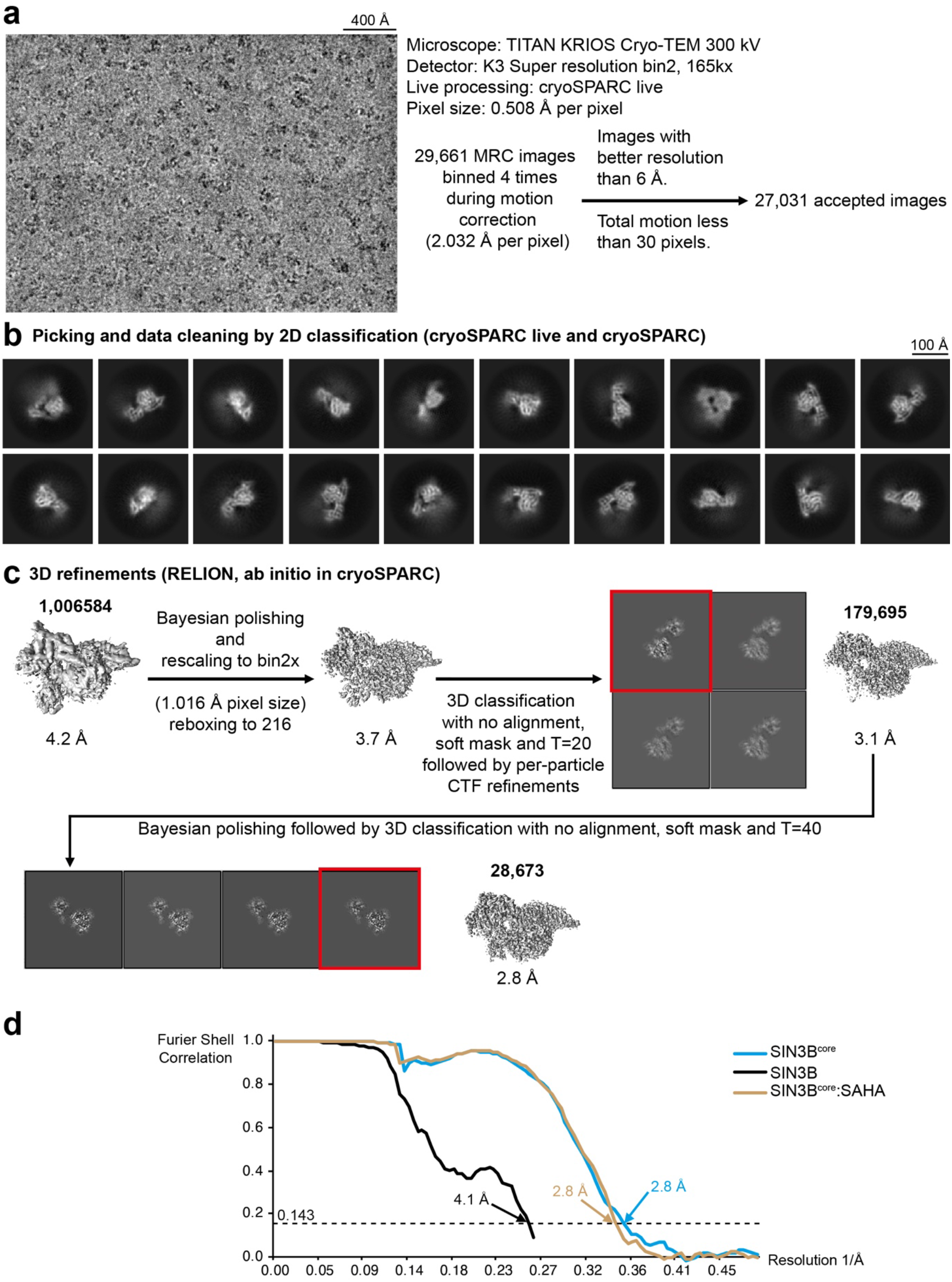
Cryo-EM analysis of the SIN3B:SAHA complex. Workflow showing a representative micrograph (low-pass filtered to 15 Å), the cryo-EM data collection parameters (**a**) and the single-particle analysis pipeline (**b-c**). N. of particles at each classification step is indicated. More details are described in the main text and in the Methods section. **d** Fourier Shell Correlation (FSC) curves for all the reconstructions are shown.

**Supplementary Fig. 5.**
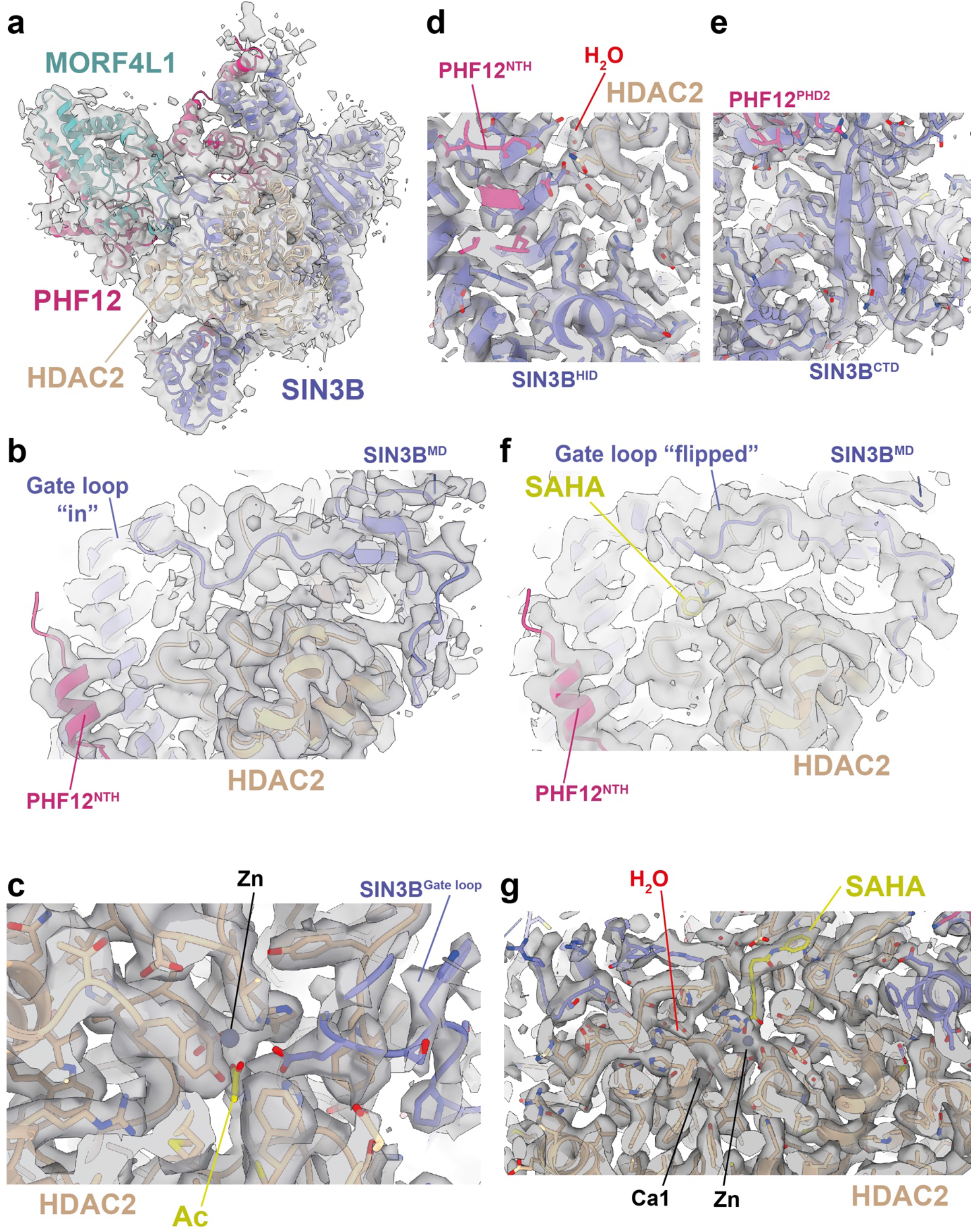
Cryo-EM analysis of the SIN3B complexes. **a-f** Gallery of snapshots showing the cryo-EM map quality for the SIN3B complex (**a**), the SIN3B^core^ complex (**b-e**), and the SIN3B:SAHA complex (**f-g**). See also Supplementary Movie 1.

**Supplementary Fig. 6.**
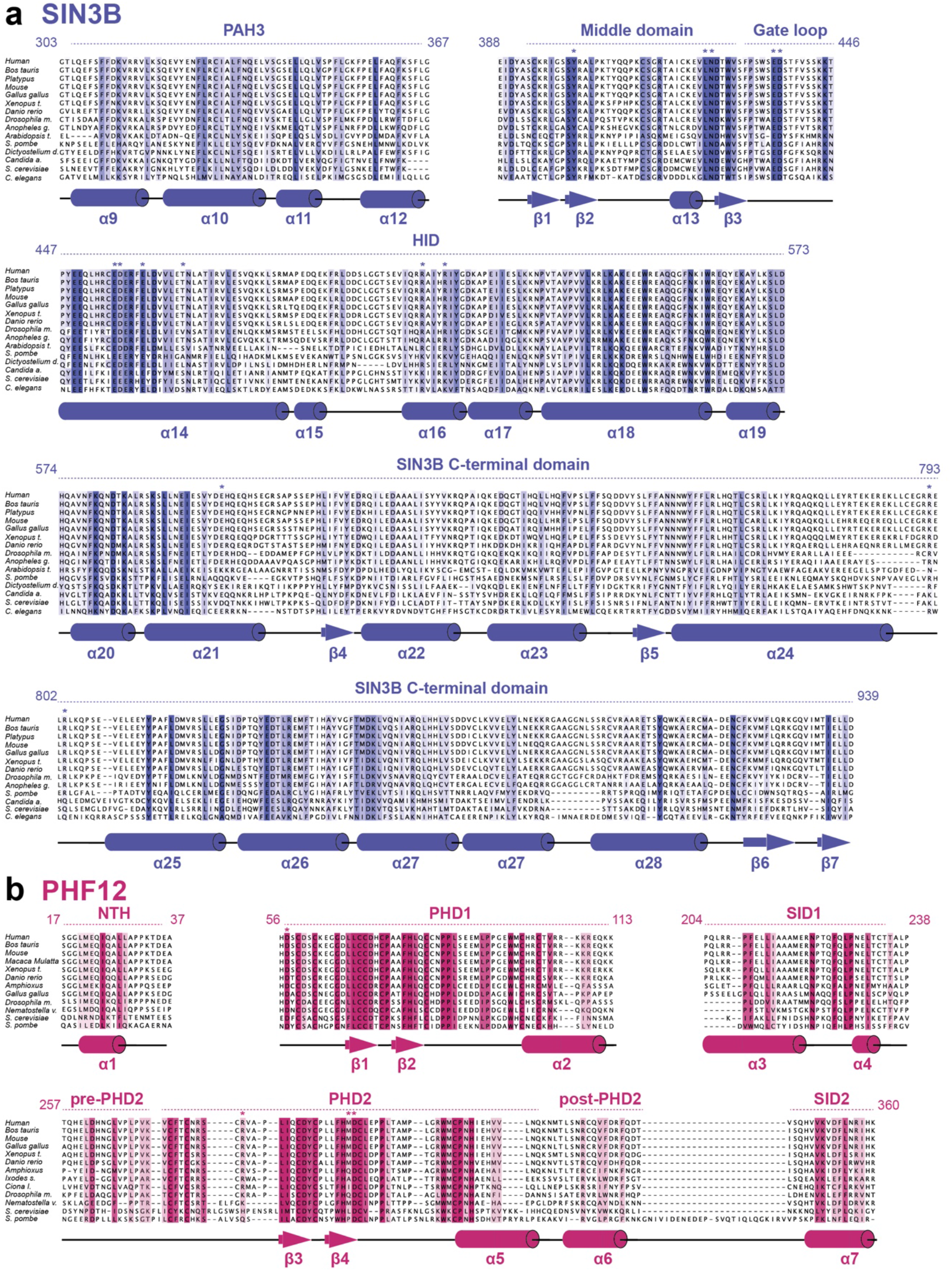
Conservation analysis on subunits of the SIN3B complex. **a-b** Sequence alignments showing conservation of region of interest within this work. Residues which are mentioned in the main text are highlighted with an asterisk.

**Supplementary Fig. 7.**
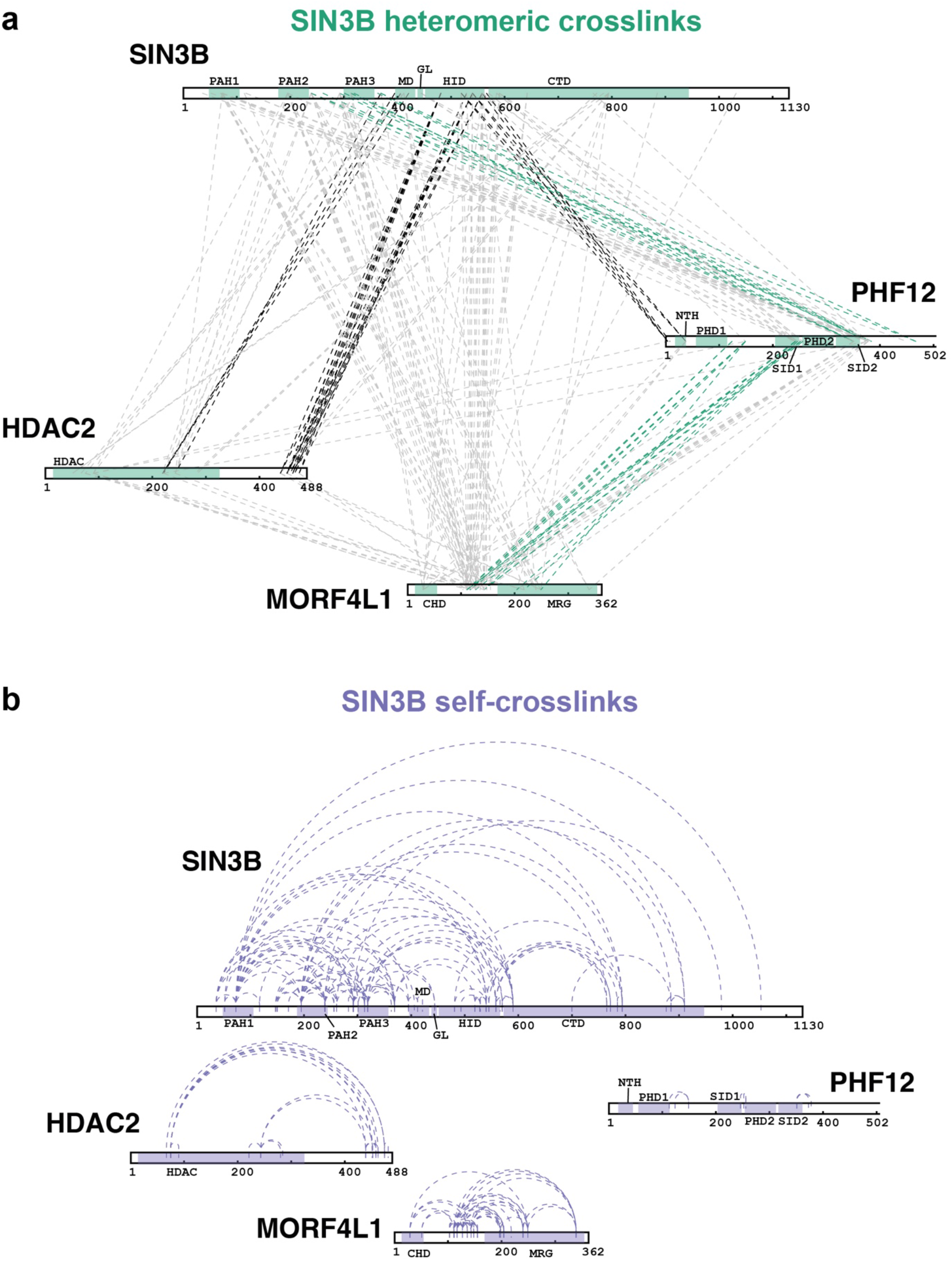
Representation of XL-MS results for the SIN3B apo complex. **a** Detected XLinked peptides are also shown on Supplementary Data 1. To obtain high confidence data for model interpretation and visualization, only the Xlinks to a value of >100 were used. Heteromeric crosslinks were visualised with xiVIEW (https://xiview.org/xiNET_website). SIN3B heteromeric crosslinks from the XL-MS data which are mentioned in the main text are highlighted in either black or green. **b** Self-crosslinks shown in the SIN3B subunits are shown.

**Supplementary Fig. 8.**
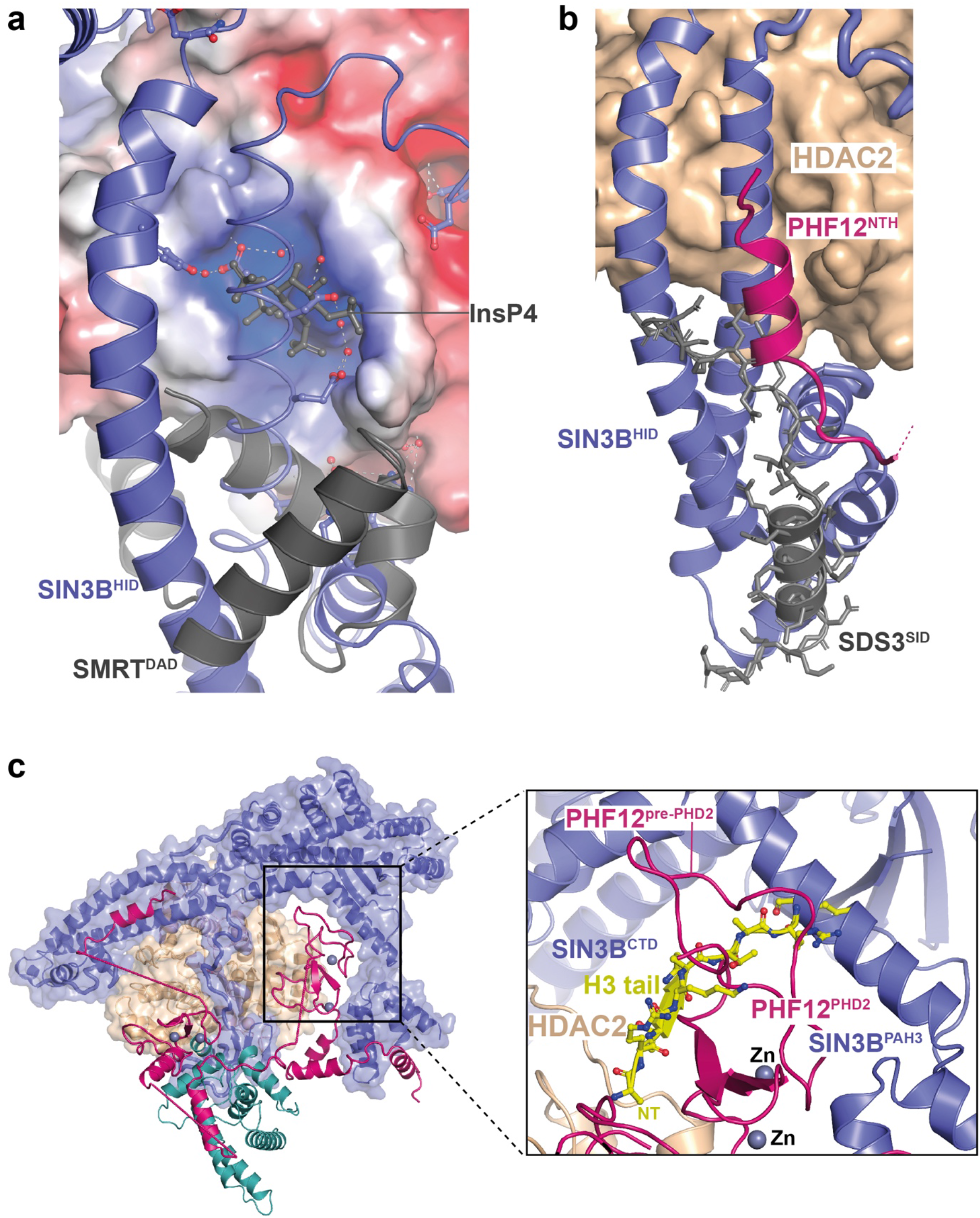
Structural comparisons of SIN3B substructures with related structures (part1). **a** Superposition of the SMRT deacetylase activation domain (DAD):HDAC3:Inositol tetraphosphate (InsP4) complex (PDB ID: 4A69) with our SIN3B complex structure (HDAC2 was the reference). **b** Superposition of the SDS^SID^:SIN3A^HID^ structure (PDB ID: 2N2H) with our SIN3B complex structure (SIN3B^HID^ as a reference). **c** Modelling of the H3 histone tail based on the structure of BHC80^PHD^:H3 (PDB ID: 2PUY) superposed to our SIN3B structure (PHD2 as a reference). This docking exercise shows severe clashes between H3 N-terminal tail and SIN3B, and PHF12.

**Supplementary Fig. 9.**
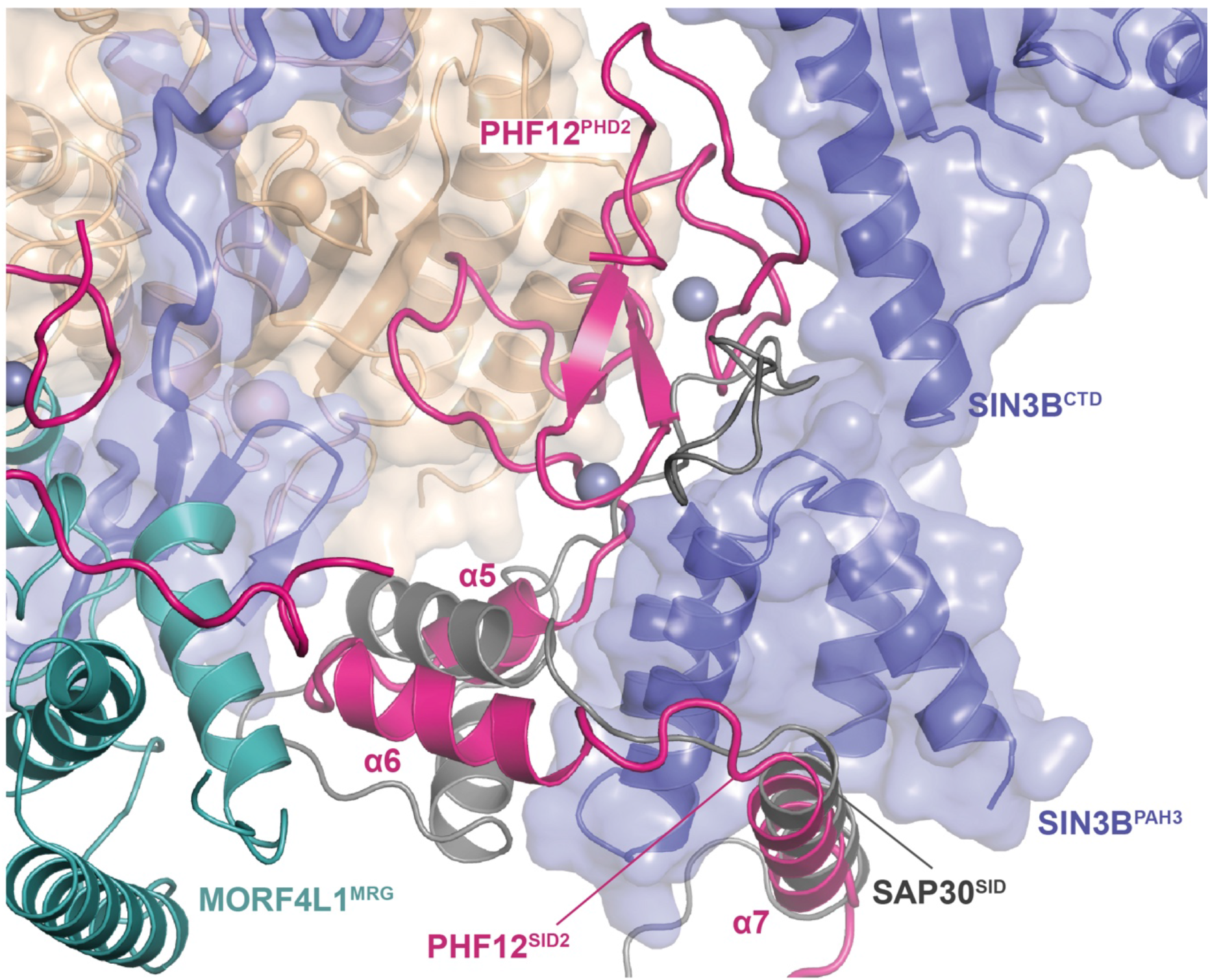
Structural comparisons of SIN3B substructures with related structures (part2). Modelling of SAP30 onto the SIN3B structure based on the structure of SAP30^SID^:SIN3A^PAH3^ structure (PDB ID: 2LD7) superposed to our SIN3B structure (SIN3B^PAH3^ as a reference).

**Supplementary Table 1.** Cryo-EM data collection, refinement and validation statistics

|  | SIN3B-FL | SIN3B | SIN3B-core | SIN3B-core:SAHA |
| --- | --- | --- | --- | --- |
| **Data collection and processing** |  |  |  |  |
| Magnification | 240,000 | 165,000 | 165,000 | 165,000 |
| Voltage (kV) | 200 | 300 | 300 | 300 |
| Electron exposure (e-/Å<sup>2</sup>) | 60 | 60 | 60 | 60 |
| Defocus range (μm) | -0.6 to -1.6 | -0.6 to -1.6 | -0.6 to -1.6 | -0.6 to -1.6 |
| Pixel size (Å) | 0.567 | 0.513 | 0.513 | 0.508 |
| Symmetry imposed | C1 | C1 | C1 | C1 |
| Initial particle images (no.) | 1,455,616 | 796,780 | 343,056 | 179,695 |
| Final particle images (no.) | 38,352 | 49,835 | 41,979 | 28,673 |
| Map resolution (Å) | 3.4 | 4.1 | 2.8 | 2.8 |
| FSC threshold | 0.143 | 0.143 | 0.143 | 0.143 |
| **Refinement** |  |  |  |  |
| Initial model used (PDB code) | 6XEC, 2LKM | 6XEC, 2LKM | 6XEC, 2LKM | 6XEC, 2LKM |
| Model resolution (Å) | 3.4 | 4.1 | 2.8 | 2.8 |
| Map sharpening B factor (Å<sup>2</sup>) | -72 | -134 | -51 | -41 |
| **Model composition** |  |  |  |  |
| Nonhydrogen atoms | 10393 | 10430 | 10442 | 10491 |
| Protein residues | 1280 | 1284 | 1284 | 1284 |
| Ligands | Zn(5), Ca(2) | Zn(5), Ca(2), ACT(1) | Zn(5), Ca(2), ACT(1) | Zn(5), Ca(2), SHH(1) |
| **B factors (Å<sup>2</sup>)** |  |  |  |  |
| Protein (min/max/mean) | 15.36/171.96/83.08 | 0.00/100.59/46.88 | 0.00/100.59/46.88 | 0.00/100.59/46.87 |
| Ligand (min/max/mean) | 93.87/216.36/142.03 | 0.00/131.51/50.19 | 0.00/131.51/50.19 | 0.00/131.51/33.84 |
| **R.m.s. deviations** |  |  |  |  |
| Bond lengths (Å) | 0.004 | 0.003 | 0.003 | 0.004 |
| Bond angles (°) | 0.783 | 0.624 | 0.624 | 0.681 |
| **Validation** |  |  |  |  |
| MolProbity score | 1.9 | 1.9 | 1.92 | 2 |
| Clashscore | 16.2 | 9.04 | 9.29 | 9.27 |
| Poor rotamers (%) | 0.7 | 2.55 | 2.46 | 2.64 |
| **Ramachandran plot** |  |  |  |  |
| Favored (%) | 96.59 | 97.47 | 97.31 | 96.91 |
| Allowed (%) | 3.41 | 2.53 | 2.69 | 3.09 |
| Disallowed (%) | 0 | 0 | 0 | 0 |

**Supplementary Movie 1. Cryo-EM maps and models of the SIN3B complex structures**. SIN3B full-length, SIN3B core, SIN3B:SAHA complex cryo-EM structures are presented.

## Data availability

The structural coordinates have been deposited in the Protein Data Bank with the following accession numbers: PDB-ID 8BPA, 8BPB and 8BPC; the EM data have been deposited in the Electron Microscopy Data Bank with the following accession numbers: EMDB-ID EMD-16147, EMD-16148 and EMD-16149.

